# Cell type classification and unsupervised morphological phenotype identification from low-res images with deep learning

**DOI:** 10.1101/533216

**Authors:** Kai Yao, Nash D. Rochman, Sean X. Sun

## Abstract

Convolutional neural networks (ConvNets) have been used for both classification and semantic segmentation of cellular images. Here we establish a method for cell type classification utilizing images taken on a benchtop microscope directly from cell culture flasks eliminating the need for a dedicated imaging platform. Significant flask-to-flask heterogeneity was discovered and overcome to support network generalization to novel data. Cell density was found to be a prominent source of heterogeneity even within the single-cell regime indicating the presence of morphological effects due to diffusion-mediated cell-cell interaction. Expert classification was poor for single-cell images and excellent for multi-cell images suggesting experts rely on the identification of characteristic phenotypes within subsets of each population and not ubiquitous identifiers. Finally we introduce Self-Label Clustering, an unsupervised clustering method relying on ConvNet feature extraction able to identify distinct morphological phenotypes within a cell type, some of which are observed to be cell density dependent.

**Author summary:** K.Y., N.D.R., and S.X.S. designed experiments and computational analysis. K.Y. performed experiments and ConvNets design/training, K.Y., N.D.R and S.X.S wrote the paper.

## Introduction

Convolutional neural networks (ConvNets) have been applied to a strikingly broad array of tasks. From stylistic transfer in artwork [1] to medical image analysis [2–5], ConvNets have been trained to successfully generate, segment, and classify images often outperforming experts across various fields. Cell biology is no exception, with groups demonstrating the segmentation [2] and classification [3–5] of single adherent cells as well as mixed populations in clusters and in suspension; however, two issues continue to prove challenging towards the application of ConvNets to single cell classification.

First, due to the small cell numbers typically obtained relative to the wide variety of accessible phenotypes, qualitative morphological differences may be observed across experiments conducted within only days of one another. These morphologies may vary greatly enough to preclude generalization of a network to novel data even when validation within the original dataset is near perfect(see Fig. 2). Second, most published methods rely on high resolution images captured through objectives with large numerical apertures (NA) or flow cytometry for suspended cells. Suspending cells utterly changes cell morphology and potentially regularizes cell shape impeding detailed classification. High NA objectives in common use have very short working distances requiring cells to be plated on thin glass coverslips before imaging. Neither case allows cells to be classified within typical thick-bottomed, plastic cell culture flasks in which cell lines are most commonly maintained.

Here we establish a method for cell type classification utilizing images taken on a benchtop microscope directly from cell-culture flasks eliminating the need for any dedicated imaging. Despite the low resolution of the images obtained and significant flask-to-flask heterogeneity, we are able to demonstrate both high accuracy and generalizability to novel data. To further measure the predictive value of the network, we recruited 27 individuals with cell culture experience to complete visual inspection and manual classification, so called “expert classification” (see Methods and Materials). Notably, when given images containing multi-cell regions, experts performed very well despite poorly classifying single cells. This finding suggests experts rely on the identification of characteristic phenotypes within subsets of each population instead of ubiquitous identifiers.

To uncover the nature and potential biological significance of these morphological phenotypes, we developed Self-Label Clustering, a novel unsupervised clustering method designed to cluster and identify distinct morphological phenotypes within one cell type. Single cells undergo significant morphological changes throughout the cell cycle, and even clonal populations span a wide range of single cell morphologies which may benefit from larger vocabulary of identifiers than those currently available. Using Self-Label Clustering, we are able to perform clustering of single cell morphologies from low-res images taken in cell culture flasks and describe two cell density-dependent phenotypes. This result highlights the use of unsupervised methods in Machine Learning to establish novel morphological identifiers as well as the observation that diffusable cell-cell communication has measurable impact on flask culture conditions.

## Results

### Single cell morphological heterogeneity

In the present study, we established an easy-to-apply pipeline for biological laboratories to classify cell types from simply microscope images. Benchtop microscope for everyday laboratory use was used for capturing cell images in flasks (Fig 1a). Representative brightfield low-res cell images were captured through the microscope and the software (Fig 1b, see Materials and Methods). The images were then maunally cropped into single cell images in squares for artificial neural network training input (Fig 1c) after properly normalizaed (Fig 1b). The ConvNet designed for cell type classification was shown in Fig 1d, with the cell image input layer, six quadruplets of convolutional, ReLU, Batch Normalization and average pooling layers, a fully connection layer and a softmax classification layer (see Materials and Methods). Once the ConvNet was given enough labeled single cell images with known cell type attributions, a successfully trained ConvNet model is expected to be capable of classifying a newly input single cell image with unknown cell type into a correct cell type with low error (Fig 1e). The basic workflow of image-based cell type classification through convolutional neural network for single adherent cell with low-res brightfield images was illustrated in Fig 1e.

**Fig 1.**
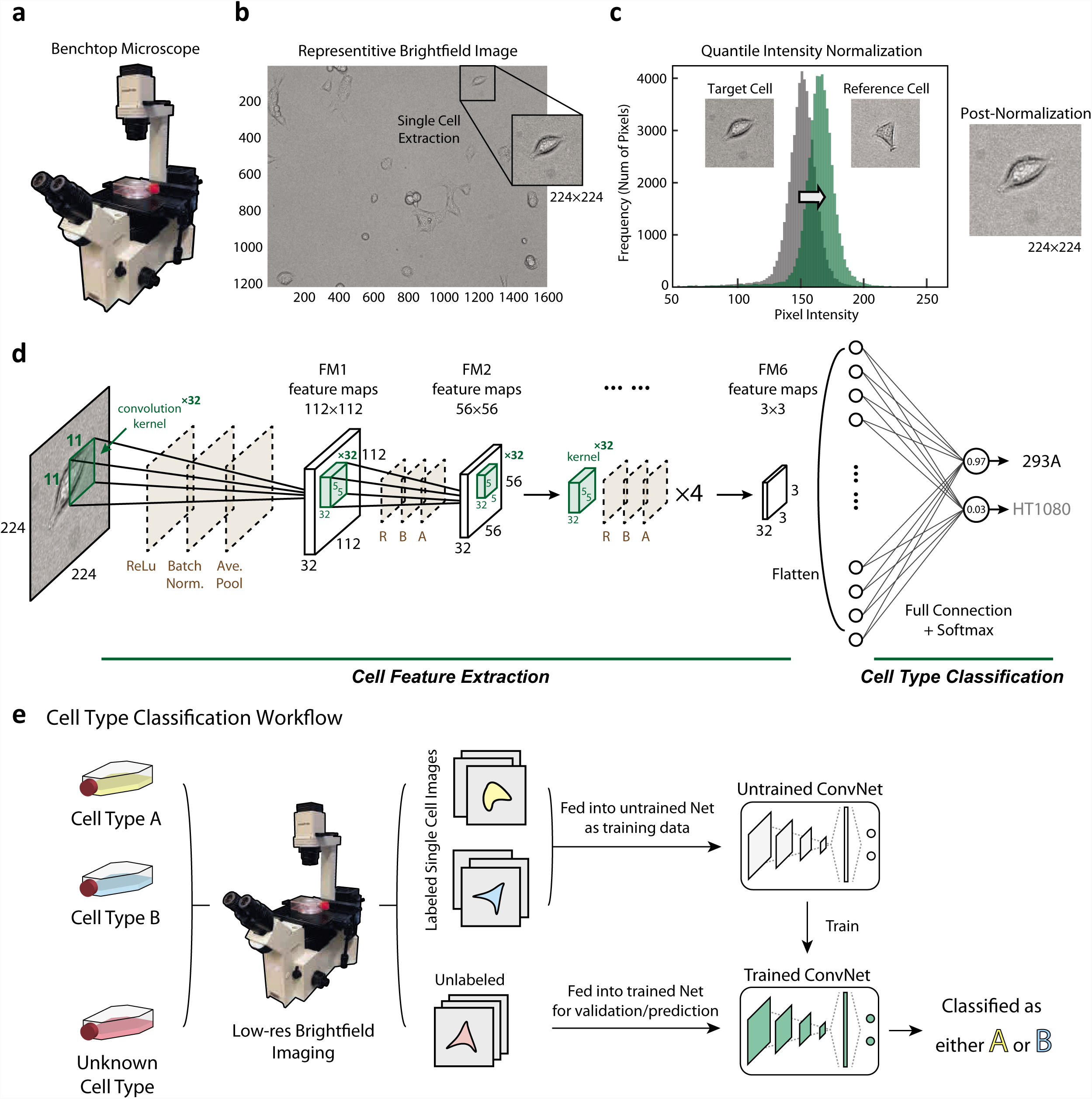
Single cell image data preparation, workflow, and artificial neural network architecture. **a.** Bench-top microscope model [info] used for capturing cell images in commonly-used, thick bottomed, plastic cell culture flasks, **b.** Example brightfield images of both multi-cell regions (1216×1616 pixels) and manually cropped single cell regions (224×224 pixels). Note the camera used was not monochromatic and the images were converted to grayscale to reduce dimensionality and improve training efficiency, **c.** Quantile normalization of the image intensity distribution was performed for each cell to a reference distribution constructed from an arbitrarily selected single cell image, **d.** Cartoon of the proposed ConvNet architecture. Six quadruplets of convolutional, ReLU, Batch Normalization and average pooling layers were constructed. A single fully connected layer was constructed before the Softmax and classification layers, **e.** Illustration of Cell Type Classification from low-res flask images through artificial neural network. Labelled cells (yellow, blue) in labelled flasks should be fed into the neural network as training data, and unlabelled flasks shoule be treated as validation data. The trained ConvNet model is able to predict cell type with low error given novel cells with unknown label.

We initially trained and validated the network on single cell images originating from a pair of flasks in a single experiment, i.e. images from one flask of cell type A (HEK-293A) were labelled positive and images from another flask of cell type B (HT1080) were labelled negative. The two flasks from one experiment are considered one ‘flask pair’ illustrated in Fig 2a (left). Randomly assigning 80% of the cells as training data and 20% as validation, for every flask pair so tested, we were able to achieve satisfactory validation accuracies in excess of 95% (Fig 2a) averaging over mini-batches, with comparable training accuracies well above expert classification with an average of 51.58% (Fig 2b). For expert classification to be a proper comparison of this training regime, all cells came from one flask pair of HEKs and HT1080s (Fig S2). The computing time spent on the ConvNet training for each trial was fairly short with generally less than 5 minutes to achieve over 95% accuracy, suggesting the method we developed is easy and fast to apply. In the mixed-population test when two cell types were mixed together and seeded overnight, the proposed ConvNet also showed excellent performance (Fig 2g).

**Fig 2.**
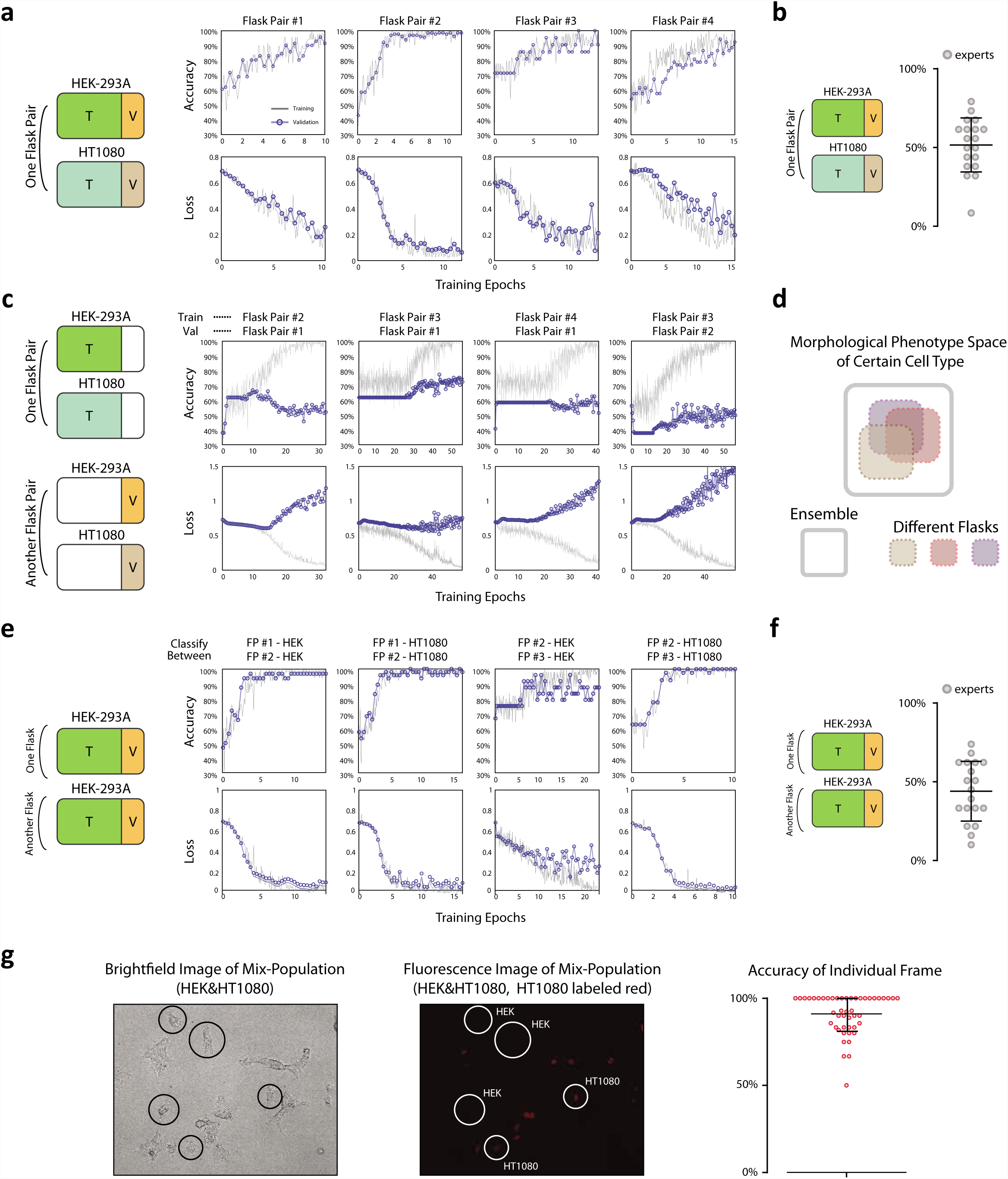
Within-flask validation shows excellent performance while cross-flask generalization is not achieved. **a.** Within-flask pair validation shows excellent model performance. Left: Illustration of training and validation dataset in this test, Right: Accurcy and loss for both training and validation across training epochs. Validation of over 95% accuracy can generally be achieved, **b.** Expert classification results corresponding to panel (a) of within-flask pair validation, **c.** Cross-flask pair validation shows poor model performance by trained neural netowrk. Left: Illustration of training and validation dataset in this test, Right: Accurcy and loss for both training and validation across neural network training epochs. Validation of over 70% accuracy can hardly be achieved, **d.** Cartoon of the proposed distribution of single cell morphological phenotypes across different flasks, **e.** Classification of the same cell type in two separate flasks shows excellent performance indicating flask-to-flask single cell heterogeneity in morphological phenotypes. Left: Illustration of training and validation dataset in this test, Right: Accurcy and loss for both training and validation across neural network training epochs. Validation of over 95% accuracy can generally be achieved, **f.** Expert classification results corresponding to panel (e) of classification between two flasks of the same cell type, **g.** Validation on mix-cell population of HEK-293A and HT1080 cells. Representative flask image was shown here with HT1080 having H2B label (red fluorescence) and HEK-293A being unlabelled fluorescently (left, mid). Accuracy of each field image was plotted in form of scatter plot (right).

We proceeded to test the generalizability of the network, selecting cells from a novel flask for the assembly of the validation set illustrated in Fig 2c (left), i.e. images from flask pair 1 of cell type A were labelled positive and added to the training set; images from flask pair 1 of cell type B were labelled negative and added to the training set; images from flask pair 2 of cell type A were labelled positive and added to the validation set; and images from flask pair 2 of cell type B were labelled negative and added to the validation set. To maintain consistency, the training options (network hyperparameters) were shared (including the training-validation ratio of 4 to 1) with the single flask pair analysis. This exercise was conducted between 4 quadruplets. Despite ideal training accuracy approaching 100% as expected, validation was poor and slightly better than random (Fig 2c, 4 pairs shown). We went on to perform the following cross-flask exercise in an effort to utilize a more robust training set.

Given 4 separate days of imaging with one flask pair imaged each day, cells from multiple flask pairs were pooled as training data with the remaining flask pairs left for validation i.e. 1, 2 and 3 flask pairs were used as training data and any flask pair of the remaining 3, 2 and 1 flask pairs respectively were used for validation. Thus this exercise yielded 12 (4choose1*3choose1), 12 (4choose2*2choose1) and 4 (4choose3*1choose1) data points respectively. Again training options were maintained. Average accuracy increased as the pool size of the training set increased, indicating day-to-day, flask-to-flask heterogeneity while great enough to prohibit generalization from a single day of imaging, does not prohibit generalization from a larger pool of training data(Fig 3e).

**Fig 3.**
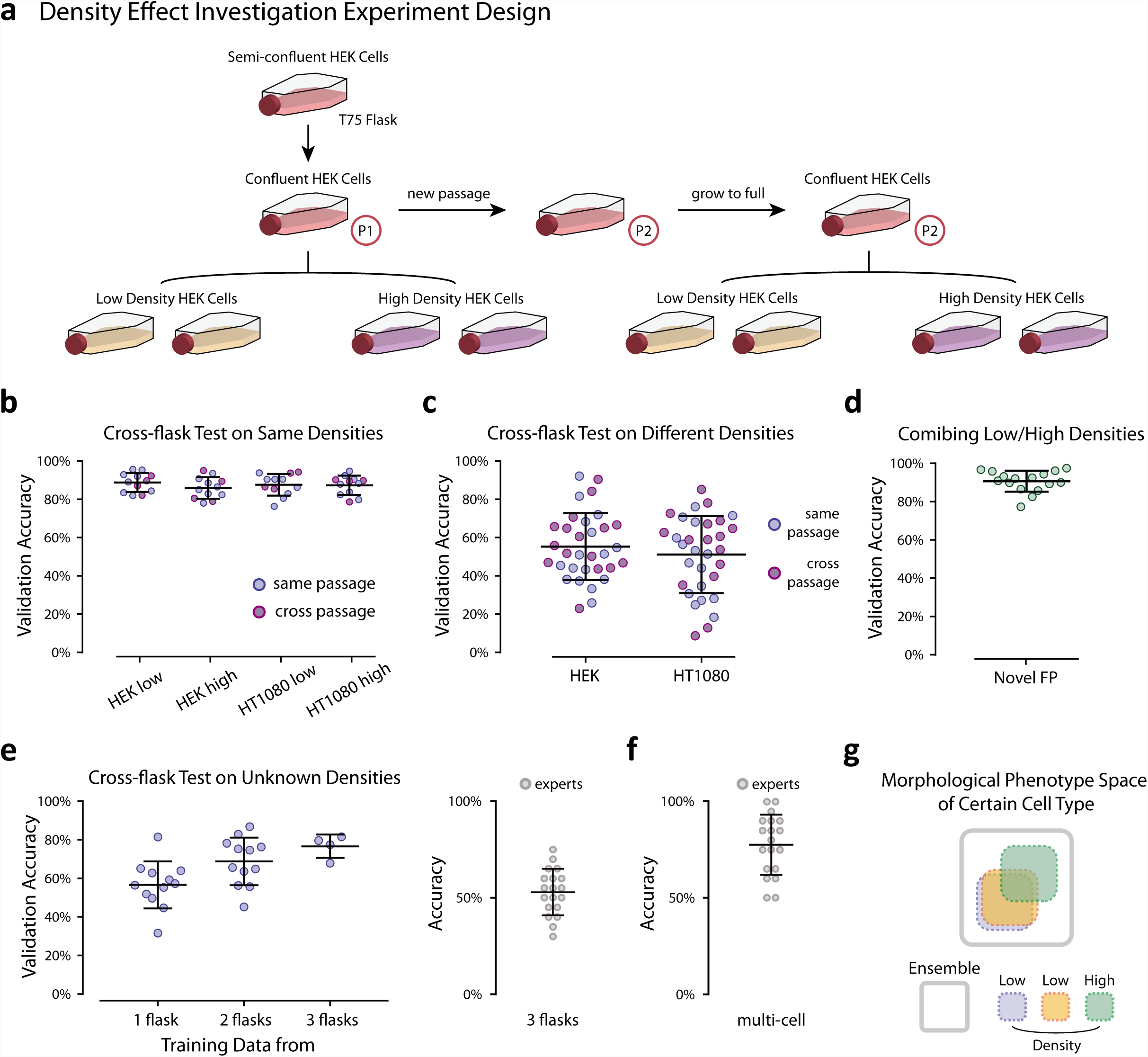
Cell density shows significant effects on single cell morphology. **a.** Cartoon of experiments designed for investigating cell density effects across flasks, **b.** Cross-flask tests with densities conserved (e.g. high density training/high density validation, low density training/low density validation). Tests for which the training and validation data are both from the same passage are labeled blue and tests for which the training and validation data are from different passages are labeled purple. There is no observed dependence on passage number, **c.** Cross-flask tests accross densities (e.g. high density training/low density validation, low density training/high density validation). Colored according to passage as in b., **d.** Training on both densities and generaing to a novel flask pair with density uncontrolled, **e.** Left: Training on pools of images from flask pairs of uncontrolled density and generating to novel flask pairs of uncontrolled density. Right: expert classification training on 3 uncontrolled flask pairs and generating to a novel flask pair of uncontrolled density, **f.** Expert classification for multi-cell frame classification of two cell types, **g.** Cartoon of the proposed distribution of single cell morphological phenotypes within a single cell type across flasks of different densities.

Given the identification of significant heterogeneity between flasks of the same cell type imaged on separate days, we went on to determine if the network was capable of accurately classifying two flasks from separate flask pairs of the same cell type illustrated in Fig 2e (left). Indeed the network was able to achieve validation accuracies well above expert classification (Fig 2e,f) performing no worse than the HEK vs HT1080 classification shown in Fig 2a. These results (Fig 2a,c,e) were all obtained from similarly prepared flasks at single cell density and imaged under the same conditions. The only unknowns are the exact number of cells seeded, imaging-day dependent biological variables (e.g. passage number), and intrinsic variability of the imaging protocol (ambeint light etc.). However seemingly negligible these unknowns, the morphological spaces occupied by the single cells from the same cell type were sufficiently different between different flasks to affect the outcome of the model learned by the network. Illustrated in Fig 2d.

### Mixing cell density achieves robust generalization

Cell density within a population is known to be a factor of affecting cellular phenotypes and cell growth through manipulating level of cell-cell interaction [6–8], and cell morphology among all cellular phenotypes can also be affected by altering cell density [9, 10]. The failure of the ConvNet to generalize between flasks for which the cell density was not controlled (Fig 2c), while all within the single-cell limit, in addition to the excellent performance of the network when classifying the same cell type from two different flasks (Fig 2e) motivated the design of experiments for which the number of cells seeded into each flask was precisely regulated for investigating single cell morphology heterogeneity under different density conditions. We went on to select two seeding densities, “high” with 0.5 million cells seeded per flask and “low” with 0.1 million cells seeded per flask (Fig 3a). Initially an intermediate number was additionally used to construct a “medium” density but the range of densities used was determined to be too narrow to warrant a subdivision (data not shown). To ensure differences in cell density were the cause of the observed variability and not another source of biological variation (e.g. passage number) or intrinsic variability of the imaging protocol, we established the following protocol (Fig 3a).

On day one, T75 cell culture medium flasks of both cell lines were prepared and incubated for 24 hrs after which time they were split into four flasks each, 2 of high density and 2 of low density. The remaining cells from each of the large flasks were seeded in a new large flask. The four density-controlled flasks were allowed to adhere for 24 hrs and were imaged within 2 hrs of one another. The 2 large flasks were maintained for one week during which time they were passaged once before the process of splitting into density-controlled flasks was repeated. In total this produced 16 flasks: 2 low density and 2 high density for each cell type imaged on two separate days from two separate passages.

Flasks of the same density were able to generalize to one another regardless of the day on which they were imaged, i.e. neither the passage number (Fig 3b) nor any other day-specific variable impedes generalization; however, networks trained on images of one density were unable to generalize to flasks of the other (Fig 3c). While these results clarified that variations in cell density, even within the single-cell regime, were the cause of this inability to generalize the network model to novel data, they highlighted a problem in the application of this, or any similar protocol, for everyday lab use. We constructed the method presented in Fig 1 to require no dedicated imaging of the user and further utilized images captured within cell culture flasks so that cell passage would not be required prior to classification; however, if only small variations in cell density preclude generalization of the model, the heterogeneity of the flasks in culture would be too great to overcome. On the other hand, networks trained on a combination of both densities were able to generalize to novel flasks of either density (Fig 3d). We went on to assemble combinations of flasks of uncontrolled cell density (discussed in the previous section) and found that, indeed, utilizing pairs or triplets of flasks for training data succeeded in increasing the validation accuracy up to 80%, well beyond expert classification (Fig 3e, right). Thus, the method presented is amenable to integration within the typical work-flow of a cell biology lab without the need for dedicated imaging or cell passage prior to data collection. The only additional requirement is that a few independent flasks of each cell type of interest should be pooled as training data to achieve better generalization.

### Expert classification vs ConvNet classification

The morphological differences between flasks of uncontrolled density were not great enough to achieve expert classification (Fig 2f). This was unsurprising considering expert performance distinguishing between two different cell types was only slightly better than random (Fig 2b); however, expert classification of images containing multiple cells was far better (Fig 3f). Images containing multi-cell regions (1216×1616 pixels) were displayed to experts in the manner described above for single cell images. In this task, experts showed decent performance achieving 77.63% accuracy on average. This result suggests experts rely on the identification of characteristic phenotypes within subsets of each population of cells instead of ubiquitous identifiers of each single cell. In other words, experts do not identify cell type based on a set of characteristics shared by the majority of cells of one cell type and absent from the majority of the other, but rather through the identification of characteristic phenotypes present in a few cells out of the group or perhaps the relative frequencies at which these minority phenotypes are observed.

### Self-Label Clustering: unsupervised morphological phenotype identification

Having established the likely presence of distinct morphological phenotypes within each cell type, we sought to determine if the network could identify these phenotypes in an unsupervised manner potentially uncovering morphological identifiers of biological significance. For this section of the work we used images of only cell type A (HEK-293A). We began by constructing “self-classes” for each cell, each class representing a series of (50) images generated out of the same cell through the augmentation outlined in Fig S1. We assigned each class a unique label and replaced the final two-class classification layer in the network architecture described in Fig 1d with an *N*-class classification layer where *N* is the number of classes determined by the number of cells in the database (Fig 4a). In this way, we constructed what we call a Self-Label ConvNet where the groups of augmentations of each cell are considered unique classes. When given each original image used to generate these classes, the trained Self-Label ConvNet model is able to return a representation of the similarities and differences among any group of the original images. These similarities and differences are in the vocabulary of novel features learned by the network training without relying on any predetermined set of morphological identifiers.

**Fig 4.**
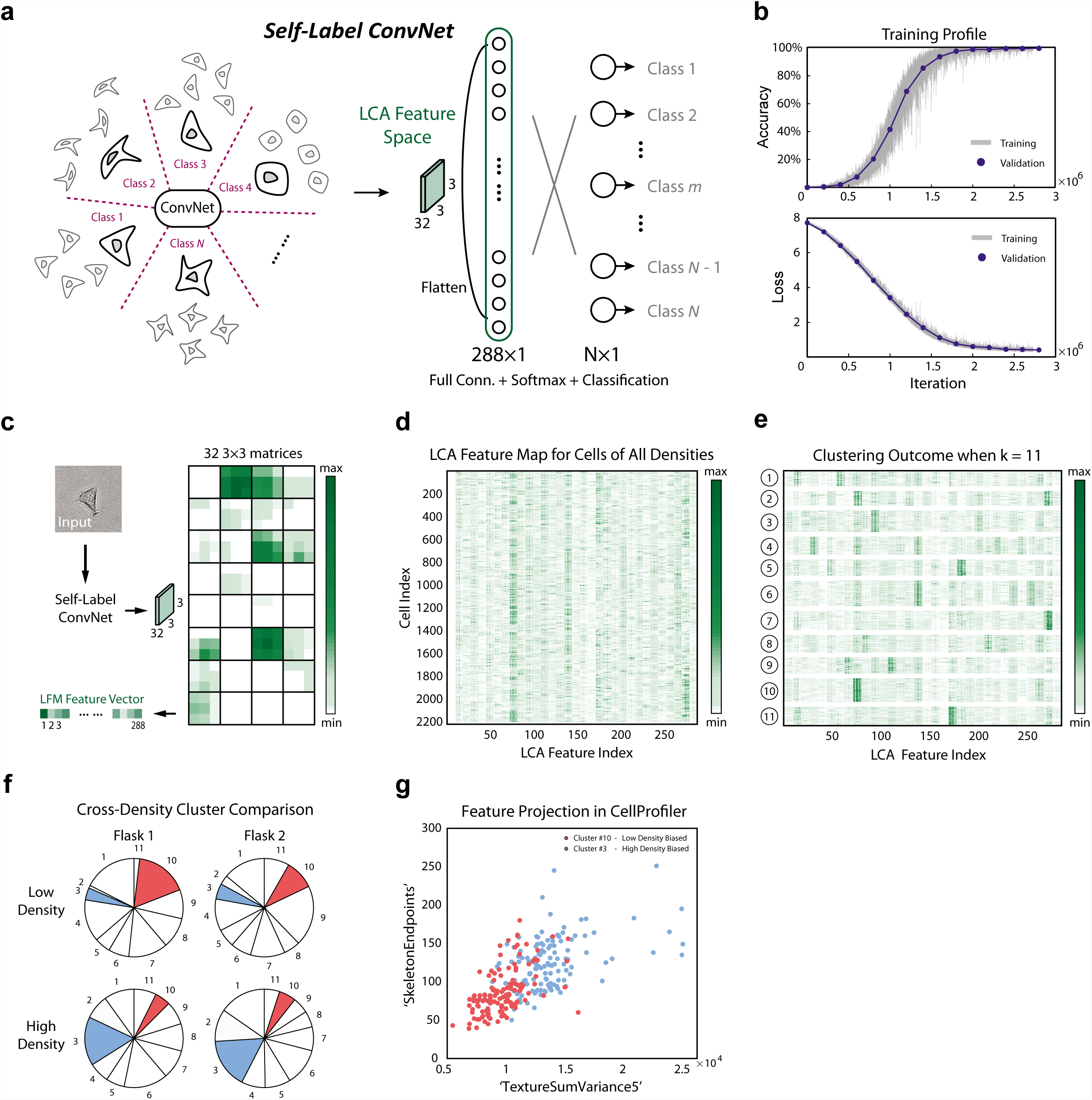
Self-Label Clustering is able to identify distinct morphological phenotypes within a single cell type. **a.** Self-Label ConvNet Architecture Illustration. The group of augemented copies for each cell are considered unique classes, yielding the same number of classes in the final layer as there are cells used to train the network. The [l]ast [c]onvolutional [a]ctivation or ‘LCA’ feature space, labeled in green, is the structure of interest for the following mophological phenotype clustering, **b.** Training profile of Self-Label ConvNet. An accuracy of nearly 100% can be achieved for both training data and validation data, and a Softmax loss of nearly 0 can be achieved for both training data and validatioan data, **c.** Workflow for acquiring the LCA feature space an example cell. Novel cells are input into the pre-trained Self-Label ConvNet and the activations of the last convolutional layer are recorded as 32 3×3 matrices for each cell input. The matrices are then flattened to a vector of length 288, each element representing one ‘feature’ of the input cell, **d.** LCA matrix: LCA feature maps for many cells across all densities (2208 cells total) are displayed as rows in a matrix (size 2208×288) with each column representing one feature in the LCA, **e.** Clustering outcome for the LCA matrix applying *k*-means to rows according to Euclidean distance with *k* = 11. Clusters are shown after reshuffling the cell indices based on their cluster index., **f.** Cross-density cluster comparison. Two flasks of two densities are shown. For each flask, the fraction of cells belonging to each of the *k* = 11 clusters are displayed. Clusters with significantly different representations between densities are colored, **g.** Morphological Analysis: Two clusters of cells dominated by low density (cluster #10) and high density (cluster #3) respectively, were analyzed. Morphological properties for cells within these clusters were calculated with CellProfiler, and two features (SkeletonEndpoints and TextureSumVariance5) were chosen to generate a 2D projection, illustrating clear distinguishability in a low dimensional morphological feature space. High density biased cluster (cluster #3) was labeled in red and low density biased cluster (cluster #10) was labeled in blue.

All cells analyzed from every density-controlled flask were included in the training data of Self-Label ConvNet in order to span the entire morphological feature space of HEK cells. We then trained the network on 80% of the data and validated on the remainder achieving close to 100% both training accuracy and validation accuracy (Fig 4b). The ‘original’ cell augmented to generate each class was excluded from both the training and validation sets. These cells were then classified through the pre-trained Self-Label ConvNet and the activations of the last convolutional layer were recorded (32 3×3 matrices). The matrices were then flattened to a vector of length 288, each element representing one ‘feature’ of the input cell(Fig 4c).

This feature space vector of size 288×1 is composed of the [l]ast [c]onvolutional layer’s [a]ctivations, which we call the “LCA Feature Space”. A matrix with rows containing LCA vectors for each cell is displayed in Fig 4d. Using features learnt from ConvNet for clustering problems has been studied previously [11] and rendered promising results. *k*-means clustering was then performed on the rows of this matrix using Euclidean distances. Unsupervised clustering of images based on their ConvNet representations has recently been shown to successfully recognize objects within the ImageNet database [11], and the Self-Label ConvNet described in this study similarly recognizes sub-classes within a single cell type without prior knowledge of these classes. Cluster numbers from 2 to 100 were tested and a peak silhouette score (Fig S4) was observed for *k* = 11 where the sum of the distances to cluster centers also appears to change slope (“elbow plot” Fig S4). The LCA matrix with rows reordered based on cluster identity is displayed in Fig 4e and the distances between pairs of LCA vectors (pairs of cells) are displayed in Fig S4 with the low distances on the diagonal indicating cells are similar to other cells belonging to the same cluster. The clusters identified through the application of the Self-Label ConvNet were then examined for the presence of any density dependence.

The fractions of the populations lying in each cluster are displayed for two low and two high density flasks in Fig 4f. The colored clusters are of special interest being relatively enriched or depleted in one density compared to the other. Approximately 16.25% of cells in high density flasks belong to cluster A (#3, red) while only 4.56% of cells at low density reside in cluster A. Similarly, approximately 13.44% of cells in low density flasks belong to cluster B (#10, green) while only 5.36% of cells at high density reside in cluster B. Cells from clusters A and B were collected and analyzed through the CellProfiler platform to determine any distinguishing morphological properties. Over 250 morphological properties were calculated for each cell in CellProfiler and used to construct a high dimensional feature space. Many dimensions in which cells from clusters A and B occupy significantly different ranges of values were found. Each dimension was then ranked by the Jenson-Shannon Distance (JSD) between the two clusters in the given dimension and the spearman correlation coefficient between the top ranking dimension and all other dimensions was calculated. In particular, we found TextureSumVariance5 a measurment of the local spatial variance (utilizing a co-occurance matrix with a scale of 5 pixels see CellProfiler Documentation) of the image to be the top ranking dimension. SkeletonEndpoints, the number of endpoints in the skeleton of the cell mask was found to be poorly correlated with TextureSumVariance5 while maintaining a high JSD. When projected onto this two dimensional space, clusters A and B are almost separable (Fig 4g). It is clear that the high density biased cluster A contains cells with a “rougher” texture and more skeleton endpoints indicating a more uneven cell boundary. Thus the application of Self-Label ConvNet revealed the presence of distinct morphological phenotypes of potential biological significance.

## Discussion

The application of ConvNets to cell type segmentation and classification promises to improve the efficiency and accuracy of image analysis in many cell biology groups. Challenges remain in improving the usability and generalizability of these methods. Protocols which do not require dedicated experiments or imaging will decrease the barrier-to-entry for groups new to these methods and we hope the utilization of images taken on benchtop microscopes directly from cell culture flasks will make the use of the techniques presented here more approachable. Morphological heterogeneity across flasks creates hurdles for network generalizability but we showed that when incorporating multiple flasks with uncontrolled density, decent generalizability is achievable.

## Materials and Methods

### Data Acquisition and Preparation

T75 cell culture medium flasks of two cell lines, HEK-293A and HT1080, were seeded with a variable number of cells within a range achieving single-cell densities following typical cell passage protocol. For cell detachment and cell passage, 0.25% tryspin (ThermoFisher Scientific) was applied to cells after washing with dPBS (ThermoFisher Scientific) twice. Cells were put in the incubator for 5 minutes to allow for detachment for cell passage. For experiments investigating density variation, low density flasks were seeded with 0.1 million cells and high density flasks were seeded with 0.5 millions cells counted by hemacytometer. Cells were allowed to adhere for 24hrs in flask before being imaged. For experiments investigating mixed-population cell culture, HEK-293A and HT1080 cells from two separate flasks were detached individually and mixed together in a 15 mL conical vial. Mixed cells were then seeded into a new flask (single cell density). The mixed-population flask was then incubated overnight for 24 hrs before being imaged. The HT1080 cell line utilized in the mixed-population studies was labelled with a fluorescent H2B tag in the nuclei in order to provide a “ground truth” comparison with the unlabelled HEK-293A cells. The mixed-population test was conducted based on the pretrained model trained with multiple controlled and uncontrolled densities.

Brightfield images (1216×1616 pixels) of cells in a typical cell culture flask (Fig 1b) were captured at 10X magnification on a standard Nikon benchtop model (Fig 1a). All tests were imaged with identical settings and preparation with the exception that in the completion of the mixed-population tests, a fluorescent image was additionally captured for each region in order to resolve the H2B nuclear label in the HT1080 population. The camera used was not monochromatic and the images were converted to grayscale to reduce dimensionality and improve training efficiency. Single cells were then manually cropped, centered within squares of 224 pixels in length. The cropped images were augmented via image rigid rotation, rigid translation, and the addition of artificial background imitating substrate irregularities before normalization (Fig S1b). For the final data preparation step, we conducted quantile normalization (Fig 1c) of the image intensity distribution to correct for any variations in light intensity across or among the fields of view. Each single cell image was normalized to a reference distribution constructed from an arbitrarily selected single cell image.

### Neural Network Design

#### Cell Type Classification ConvNet

A graphical representation of the cell type classification neural network designed in the present work via MATLAB 2018a (MathWorks) inspired by the structure of AlexNet [1] was displayed in Fig 1d. The convolutional neural network is composed of the image input layer of size 224×224×1, followed by six quadruplets of a convolutional layer, a rectified linear unit layer, a batch normalization layer, and an average pooling layer, as well as one fully connected and one softmax layer at the end. The first convolutional layer has 32 kernels each of size 11×11 pixels and the subsequent five each have 32×32 pixel kernels of size 5×5 pixels. Stochastic gradient descent [1] was chosen as the learning algorithm and the learning rate was set to be fixed at 10^−4^. We observed a learning rate within 10^−3^ to 10^−4^ to be appropriate for the tests conducted in this work, and the application of learning rate decay did not have a meaningful impact. Mini-batch size was set to be approximately 5% of the size of the dataset in each test and the mini-batches were designed to be shuffled randomly every epoch throughout the training process to enhance model validation performance. The ConvNet training was performed utilizing GPU (NVIDIA).

#### Self-Label ConvNet

A graphical representation of the Self-Label ConvNet designed for cell morphological phenotype clustering within one cell type via MATLAB 2018a (MathWorks) was displayed in Fig 4a. The number of cells in the ensemble was indicated by *N* (in this study *N* = 2208). *N* classes were constructed in Self-Label ConvNet in the final layer (Softmax classification) instead of two classes for the cell type classification, while other layers before the final layer remained exactly unchanged from Fig 1d, the cell type classification ConNet. Each class in Self-Label ConvNet represents the combination of a series of *m* images (in this study *m* = 50) generated out of the same cell image through the augmentation outlined in Fig S1, and each class was then assigned a unique label of (”1”,”2”, through “*N* “) indicating *N* categories fo distinguished morphological phenotypes throughout the ensemble. The training data of Self-Label ConvNet was then composed of *N × m* single cell images, leading to a much heavier computational cost for neural network training with around 3 million iterations to achieve stable accuracy and loss (Fig 2b). Once the Self-Label ConvNet was successfully trained to a near 100% accuracy, the activations of the last convolutional layer of the ConvNet were investigated (see Results, Fig 4c,d).

### Expert Classification

To evaluate neural network performance and additionally to investigate similarities/contrasts between human and network feature identification, an expert classification survey was distributed to 20 individuals (Fig S). Four parts were included in the survey: 1. Within-flask pair classification between HT1080 and HEK-293A cells (Fig 2b), 2. Classification between two flasks of a single cell type (cross-flask pair identification) (Fig 2f), 3. Classification of two cell types when including multiple flask-pairs with uncontrolled density as training data and a novel flask-pair as validation data(Fig 3e), and 4. multi-cell frame classification of two cell types in a single flask pair (Fig 3g). Illustrated in Fig S3, experts were given 40 labeled images of cell type A and 40 labeled images of cell type B as preparation on one side of the survey paper. They were then asked to classify 20 new unlabelled cells as either type A or type B on the back side of the survey paper in parts 1 through 3. In part 4, 40 multi-cell images were given for each cell type as labeled data and 20 new images were given as unlabelled data for expert classification. The performance of expert classification tasks were evaluated by the distribution of accuracy achieved by each individual expert and also average accuracy and standard deviation of all experts’ performances shown in Fig 2b,f and Fig 3e,f for the 4 designed tasks respectively.

### CellProfiler

CellProfiler is a free, open-source image anlysis software aiding users in image segmentation and the calculation of morphological features [12, 13] as well as subsequent statistical inference through CellProfiler Analyst [14]. CellProfiler was utilized in this study to provide connections between a standard morphological feature space constructed using a rigorous, well established pipeline and the clusters generated by the Self-Label ConvNet defined only by the novel, uncharacterized features learned by the neural network. To establish the morphological features which differ between the clusters, we calculated the Jensen-Shannon Distance between the two clusters in each dimension. The Jensen-Shannon Distance (JSD) is the square root of the Jensen-Shannon Divergence, also known as the total divergence to the average or the information radius (IRad), which itself is based on the Kullback-Leibler Divergence. The JSD between two (or more) probability distributions is a measure of the dissimilarity of those distributions. For two distributions, and using the natural logarithm, the JSD is bounded by 0 for identical distributions and ln(2) for dissimilar distributions. We found the clusters generated by the Self-Label ConvNet to have high JSD for many features calculated with CellProfiler. To select two dimensions on which to project the clusters to display their separability via morphological features, we went on to calculate the spearman correlation between each dimension and the dimension with the highest JSD (TextureSumVariance5). We then selected the second dimension (SkeletonEndpoints) to have high JSD but low correlation (See SMX). Alternatively, pairs of dimensions may be more rigorously scored by measuring the separability of the clusters through a Support Vector Machine. We considered this method satisfactory for construction of the illustrative panel, Fig 4g.

### Code Availability

All codes in the present work including but not limited to data preparation (singel cell cropping, image augmentation and image normalization), ConvNet model construction, Self-Label Clustering package can be found at https://github.com/sxslabjhu/CellTypeClassification.

